# Jointly modeling deep mutational scans identifies shifted mutational effects among SARS-CoV-2 spike homologs

**DOI:** 10.1101/2023.07.31.551037

**Authors:** Hugh K. Haddox, Jared G. Galloway, Bernadeta Dadonaite, Jesse D. Bloom, Frederick A. Matsen, William S. DeWitt

**Author notes:** To whom correspondence should be addressed. (H.K.H.), (J.D.B.), (F.A.M.), or (W.S.D.). H.K.H. (Author One) contributed equally to this work with J.G.G. (Author Two). Conceptualization, J.D.B., H.K.H., W.S.D., J.G.G., F.A.M.; experiments, B.D.; computational methods, J.G.G., H.K.H., W.S.D., F.A.M., J.D.B.; computational analysis, J.G.G., H.K.H., W.S.D.; writing—original draft, H.K.H., J.G.G., W.S.D.; writing—review & editing, H.K.H., J.G.G., W.S.D., F.A.M., J.D.B., B.D.; supervision, H.K.H., W.S.D., F.A.M., J.D.B.; funding acquisition, F.A.M., J.D.B. J.D.B. is on the scientific advisory boards of Apriori Bio, Aerium Therapuetics, Invivyd, and the Vaccine Company. J.D.B. consults for Pfizer and GSK. J.D.B. and B.D. are inventors on Fred Hutch licensed patents related to the deep mutational scanning of viral proteins.

## Abstract

Deep mutational scanning (DMS) is a high-throughput experimental technique that measures the effects of thousands of mutations to a protein. These experiments can be performed on multiple homologs of a protein or on the same protein selected under multiple conditions. It is often of biological interest to identify mutations with shifted effects across homologs or conditions. However, it is challenging to determine if observed shifts arise from biological signal or experimental noise. Here, we describe a method for jointly inferring mutational effects across multiple DMS experiments while also identifying mutations that have shifted in their effects among experiments. A key aspect of our method is to regularize the inferred shifts, so that they are nonzero only when strongly supported by the data. We apply this method to DMS experiments that measure how mutations to spike proteins from SARS-CoV-2 variants (Delta, Omicron BA.1, and Omicron BA.2) affect cell entry. Most mutational effects are conserved between these spike homologs, but a fraction have markedly shifted. We experimentally validate a subset of the mutations inferred to have shifted effects, and confirm differences of >1,000-fold in the impact of the same mutation on spike-mediated viral infection across spikes from different SARS-CoV-2 variants. Overall, our work establishes a general approach for comparing sets of DMS experiments to identify biologically important shifts in mutational effects.

**Significance Statement:** Amino-acid mutations to a protein have effects that can shift as the protein evolves or is put under new selective pressure. The effects of amino-acid mutations to a specific protein under a defined selective pressure can be measured by deep mutational scanning experiments. Here, we devise an approach to quantify shifts in mutational effects between experiments performed on different homologs (i.e. variants) of the same protein, or on the same protein selected under different conditions. We use this approach to compare experiments performed on three homologs of SARS-CoV-2 spike, identifying mutations that have shifted in their effect on spike-mediated viral infection by >1,000 fold across SARS-CoV-2 variants.

---

Deep mutational scanning (DMS) is a high-throughput experiment that measures the effects of thousands of mutations to a protein (1, 2). It has been used to study a wide variety of proteins, helping to map how mutations affect phenotypes such as binding, catalysis, stability, and viral replication, among others (1–11). An additional application of DMS is to perform it on different homologs of the same protein (12–19), or on the same homolog under different selective conditions (20–27). In such cases, comparing the results can reveal how much the effects of specific mutations have shifted between homologs (due to epistasis) or between conditions (due to distinct selective pressures).

However, when comparing DMS experiments, a key challenge is determining whether observed differences in mutational effects are due to real biological signal or the noise inherent in any high-throughput experiment. Previous studies have addressed this challenge by separately inferring mutational effects in each experiment, and then trying to identify mutations with differences between experiments that are significantly larger than the experimental noise (12). However, this approach does not consider certain aspects of the data that are informative. First, most mutations have similar effects across protein homologs or selective pressures, with only a small fraction of mutations typically having large shifts in their effects (12–17, 20–25, 28). Second, differences in measured mutational effects across homologs or conditions are more likely to represent true biological shifts than noise when the measurements for a mutation have higher confidence (such as when the mutation is present in more unique variants in the experimental libraries). If one infers mutational effects for each experiment independently, this fails to directly use these two features of the data when assessing whether the differences represent real shifts or noise.

Here, we present an approach that jointly infers mutational effects across multiple experiments, and also assesses how much the effect of each mutation has shifted across homologs or conditions. As part of this approach, the inferred shifts in effects are regularized, encouraging their values to be zero unless nonzero shift values are strongly supported by the data. Therefore, our approach effectively allows all experiments to inform a shared set of mutational effects, while also allowing a subset of these effects to be shifted across homologs or conditions when the data strongly support it. Our statistical methods apply sparse estimation techniques, a family of methods—most notably the *lasso*—for inferring compressed or structured models (29).

We implement this approach in an open-source Python package called `multidms` and use it to compare DMS experiments of three homologs of SARS-CoV-2 spike. We find that most mutations have similar effects among the homologs, but that a few have large shifts in their effects. The sites with large shifts in mutational effects span several regions of spike, including the N-terminal domain (NTD), receptor-binding domain (RBD), and regions involved in conformational dynamics and inter-protomer packing. These sites tend to cluster near each other in spike’s 3D structure, but are often far away from non-identical sites that differ in amino-acid sequence between homologs, suggesting that many shifts are due to long-range epistasis. We experimentally validate a subset of the inferred mutational shifts, identifying some mutations that differ in their effect on spike-mediated viral entry by *>*1,000-fold between homologs.

## Results

### Model summary

We assume the DMS data report an experimentally measured functional score for each protein variant from each homolog or condition, where a variant corresponds to a unique protein sequence covering the entire mutagenized region. This requirement is usually satisfied by DMS experiments that either sequence the entire length of the mutagenized gene, or use barcoding to link full gene sequences to barcodes (8, 30). Each mutation may be present in one or more variants, potentially in combination with other mutations.

Given data from multiple DMS experiments, we devised a custom version of a global-epistasis model (31) to infer mutational effects in one of the experiments — defined as a *reference experiment* — and *shifts* in mutational effects in each of the other experiments relative to the reference (Figure 1). We present our model with formal notation in *Materials and Methods*, and summarize it informally here. Each variant *v* from each experiment *d* is modeled to have a *latent phenotype*:

**Fig. 1.**
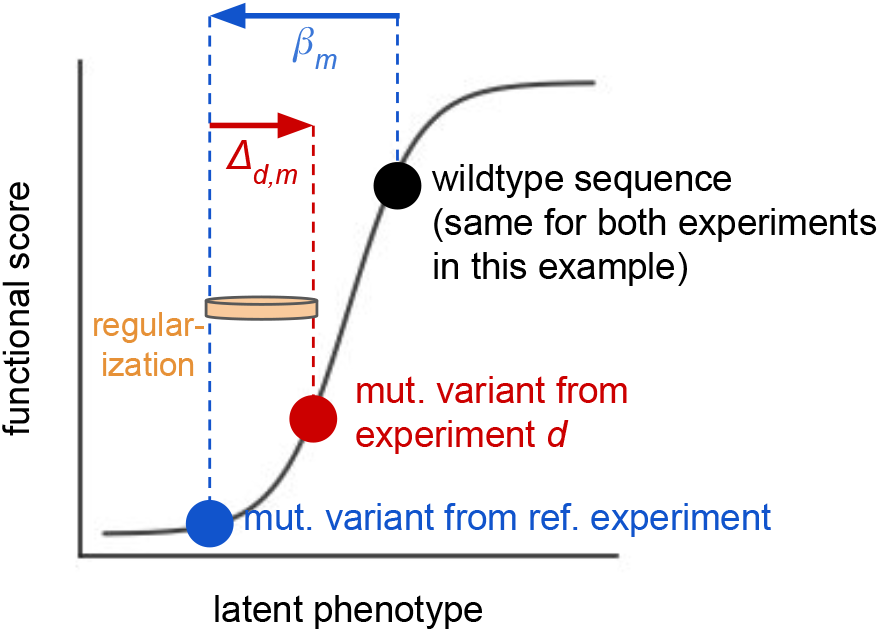
Approach to model multiple DMS experiments with a single global-epistasis model. One experiment is chosen to be a reference experiment, and the wildtype sequence from that experiment (black dot) has an inferred latent phenotype. The phenotypes of all variants from all experiments are defined relative to this wildtype sequence. Mutations to the wildtype sequence change its latent phenotype in an additive fashion. For each mutation *m*, the model infers the latent effect of *m* in the reference experiment as *β*_*m*_. For each non-reference experiment *d*, it also infers the shift in the mutation’s effect in *d* relative to the reference experiment as ∆_*d,m*_. Lasso regularization is used to drive inferred shifts ∆_*d,m*_ to zero unless they are strongly supported by the data, as symbolized by the tan rubber band. The model also infers a global-epistasis function (grey curve), which maps changes in latent phenotype to predicted changes in functional score. The blue dot shows the inferred location of a variant from the reference experiment with a single mutation *m* with effect *β*_*m*_. The red dot shows the inferred location of the same mutant variant from a non-reference experiment *d*. These variants have different predicted functional scores, which in the model is due to the inferred shift ∆_*d,m*_. If this nonzero shift greatly improves model fit it will be resistant to regularization. This example assumes that the two experiments have the same wildtype sequence and differ in the selection conditions; the situation is slightly more complicated if the two experiments differ in the wildtype sequence of each homolog, which is described in the main text.

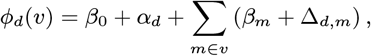

where *β*_0_ is the latent phenotype of the wildtype (i.e., unmutated) sequence from the reference experiment, *α*_*d*_ is an experiment-specific offset parameter described below, *β*_*m*_ is the latent effect of mutation *m* in the reference experiment, and Δ_*d,m*_ is the shift in the mutation’s latent effect in experiment *d* relative to the reference experiment. We fix ∆_*d,m*_ = *α*_*d*_ = 0 when *d* is the reference experiment. The summation term adds the effects of all mutations that separate *v*’s sequence from the reference experiment’s wildtype sequence. Thus, all variants from all experiments are modeled relative to the reference experiment’s wildtype sequence. If the experiments have different wildtype sequences (i.e., were performed on different homologs of a protein), then the model treats wildtype sequences from non-reference experiments as variants with mutations, and models them based on the effect of each mutation separating the homologs. The same logic is used for mutant variants from such non-reference experiments. If one or more of the mutations separating homologs are not included in the DMS libraries (e.g., indel mutations), then the individual *β*_*m*_ and ∆_*d,m*_ parameters for these mutations cannot be inferred from the data, in which case we remove them from the summation term and use a single parameter *α*_*d*_ to infer their combined effect. This parameter can be fixed to zero if all relevant mutations have DMS measurements. The *SI Appendix* describes the above logic in greater detail.

Next, the model uses a *global-epistasis function* (31) to map latent phenotypes to predicted functional scores:

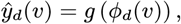

where *g* is a monotonically increasing nonlinear function, such as the sigmoid in Figure 1, which allows mutations to have nonadditive effects on functional scores, and helps to model saturation effects from global (i.e. nonspecific) epistasis or experimental limits of detection (31–35). Previous studies have explored a variety of functions for mapping latent phenotypes to functional scores (11, 25, 31, 36–38), ranging from functions that are more flexible (e.g., splines) to ones that are more constrained (e.g., sigmoids), and `multidms` allows the user to select among various options for *g*. In this study, we used the following sigmoidal function:

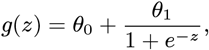

where the *θ* parameters allow the sigmoid to be fit to the range of observed functional scores. We chose a sigmoid since more flexible functions did not improve model fit (data not shown).

To estimate the above model parameters, we minimize an objective that combines a loss term (measuring the difference between predicted and experimentally measured functional scores), and a lasso regularization term that encourages the inferred shifts to be zero. Specifically, for each shift parameter, the lasso term adds a penalty that scales linearly with the absolute value of the shift. If a shift in the effect of a specific mutation is strongly supported by the experimental data, then inferring a nonzero shift for this mutation will decrease the loss term enough to overcome the regularizing effect of the lasso term. How strongly a shift is supported by the data is influenced by multiple factors, including whether the shift substantially minimizes the loss term for an individual variant, an example of which is illustrated in Figure 1, and whether it does so repeatedly across many unique variants. The strength of the lasso penalty can be tuned so that it is strong enough to drive shifts to zero if they are only weakly supported by the data, helping reduce the impact of experimental noise, but not so strong that it prevents the model from learning authentic signal.

We implemented the model described above in a Python package called `multidms`. See https://github.com/matsengrp/ multidms for the code; see https://matsengrp.github.io/multidms/ for the documentation.

### Inferring shifts in mutational effects between SARS-CoV-2 spike homologs

We applied the above approach to infer shifts in mutational effects between three homologs of the SARS-CoV-2 spike protein: Delta, Omicron BA.1, and Omicron BA.2. The Delta and BA.1 homologs are separated by 43 amino-acid mutations and indels (97% identity), while the two Omicron homologs are separated by 27 mutations (98% identity). Given the high percent identity, we expected most mutations to have similar effects among homologs.

As input to our analysis, we used previously published DMS data on how mutations to the spikes of Delta and BA.1 affect spike-mediated viral entry in the context of pseudotyped lentiviruses (39), as well as comparable data for the spike of BA.2 that we generated in new experiments performed for the current study. The DMS experiments used spike mutant libraries that each contained ∼66,000 to 139,000 variants, with an average of ∼2 to 3 amino-acid mutations per variant (Figure S1A), and with each amino-acid mutation seen in an average of ∼10 variants (Figure S1B). As described in (39), these libraries were designed to largely include only amino-acid mutations that are observed among the millions of sequenced natural SARS-CoV-2 isolates, which excludes many highly deleterious mutations. Functional scores were calculated based on the ability of each spike variant to mediate pseudovirus infection of cells expressing ACE2, as described in (39), and then truncated at a common lower bound across all experiments based on the dynamic range of the assay (see *Materials and Methods*). For each homolog, the DMS experiment was performed with at least two biological replicates starting from independently generated libraries. The functional scores were only moderately correlated among variants that were present in both replicate libraries for a given homolog (Pearson *R ∼*0.5-0.9; Figure S2A), indicating a non-trivial level of noise in the data.

We fit a single `multidms` model for the three homologs, using just one of the DMS experiments for each homolog. We used BA.1 as the reference because it had the lowest level of noise (Figure S2A), but found that the results correlated well between choices of reference (Figure S3). In fitting the model, we tested a wide range of lasso penalty strengths, choosing one that was strong enough to reduce signs of overfitting, but not so strong that it prevented the model from learning apparent signal in the data (see *Materials and Methods*; Figure S4). To gauge reproducibility, we repeated the entire fitting procedure on a separate set of replicate DMS experiments, using one experiment per homolog as above.

### Most mutational effects are conserved between homologs, but a subset have large shifts

We analyzed the inferred mutational effects, focusing on the 5,934 mutations seen at least once across all three homolog DMS experiments. Because we used BA.1 as the reference, the inferred *β*_*m*_ parameters quantify the effects of mutations in BA.1, while the inferred shift parameters quantify shifts in effects in Delta or BA.2 relative to BA.1.

Figure 2A shows the distribution of inferred mutational effects in BA.1, averaged between the two models independently fit to different replicate datasets. Nearly all mutations to stop codons had strongly deleterious effects, indicating they greatly impaired spike-mediated viral entry as expected. Most nonsynonymous and in-frame codon-deletion mutations had roughly neutral effects, while a subset had deleterious effects. These patterns are expected given that the library-design strategy targeted amino-acid mutations observed in natural SARS-CoV-2 sequences, most of which are expected to be well tolerated.

**Fig. 2.**
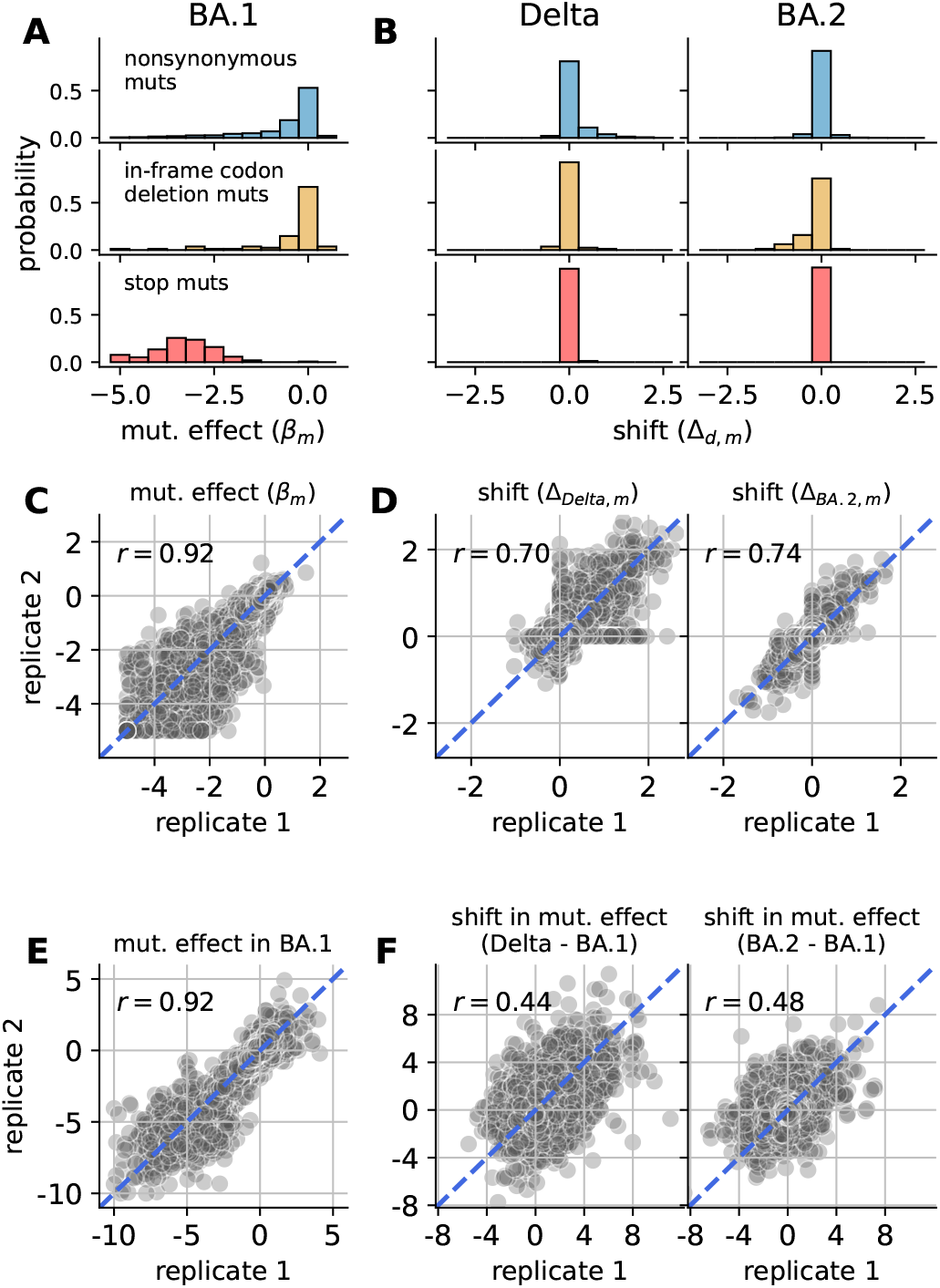
Inferred mutational effects. **(A)** Distribution of inferred mutational effects in BA.1 (*β*_*m*_) averaged between replicates and partitioned by categories: nonsynonymous mutations, in-frame codon-deletion mutations, and mutations to stop codons. In panels A and C, *β*_*m*_ values are clipped at a lower limit of -5. **(B)** Corresponding distributions of inferred shifts in mutational effects (∆*d,m*) for Delta or BA.2 relative to BA.1, averaged between replicates. **(C)** Correlation of mutational effects in BA.1 between replicates. *r* reports the Pearson correlation coefficient. **(D)** Correlation of shifts in mutational effects for Delta (left) or BA.2 (right) between replicates. **(E)** and **(F)** are similar to C and D, but show results from separately fitting a single model to each homolog’s DMS experiment, instead of the joint-fitting approach. Panel E shows the correlation of mutational effects inferred from replicate BA.1 experiments, clipped at a lower limit of -10. Panel F shows the correlation of shifts for either Delta or BA.2 relative to BA.1 as inferred from a given set of replicate experiments, where shifts are computed by subtracting the mutational effect inferred for either Delta or BA.2 by the mutational effect inferred for BA.1.

Figure 2B shows the distribution of inferred shift parameters for either Delta or BA.2 relative to BA.1, also averaged between the independent fits to different replicates. Nearly all mutations to stop codons had shifts of zero, which is expected since these mutations should be equally deleterious in each homolog. Most nonsynonymous and codon-deletion mutations also had small shifts near zero, indicating that most mutational effects are conserved between homologs. However, a small fraction of mutations had large shifts in effects between BA.1 and Delta or BA.2 (Figure S5), suggesting that the effects of these mutations are influenced by strong epistatic interactions.

Both *β*_*m*_ and shift parameters were well correlated between the independent fits to different replicates (*R ∼*0.7 for shift parameters; Figure 2C and D), showing that estimates were reproducible for the entire experimental/computational workflow. For comparison with the joint model, we separately fit a single global-epistasis model to each homolog’s DMS data and then computed shifts by subtracting the inferred mutational effects between homologs. While mutational effects inferred for BA.1 were still well correlated between replicate fits (Figure 2E), the inferred shifts had a much lower correlation (*R ∼*0.4-0.5; Figure 2F), showing that the regularized shifts inferred by the joint model were much more reproducible across noisy experiments, and so more likely to reflect real biological signal.

### Shifted mutations occur in multiple domains of spike

Sites with shifted mutational effects occurred across the length of spike (Figure 3). Several trends were apparent. If a mutation was strongly shifted, then at least a few other mutations at the same site or neighboring sites also tended to be shifted in the same direction, which could indicate a common epistatic mechanism underlying the group of shifts. This clustering of mutations with shifted effects results in punctuated patterns of shifts across primary sequence. The shifts in Delta were mostly positive, as further discussed below. BA.2 had a mix of positive and negative shifts, with negative shifts concentrated in the NTD and positive shifts concentrated in other domains. Many shifts were unique to either Delta or BA.2, though some shifts were similar in both (e.g., sites 568-572; Figure S6), suggesting that some mutations were uniquely shifted in BA.1 relative to both Delta and BA.2. See https://matsengrp.github.io/SARS-CoV-2_spike_multidms/spike-analysis.html#shifted-mutations-interactive-altair-chart for an interactive heat map that enables more detailed analyses of the mutational shifts.

**Fig. 3.**
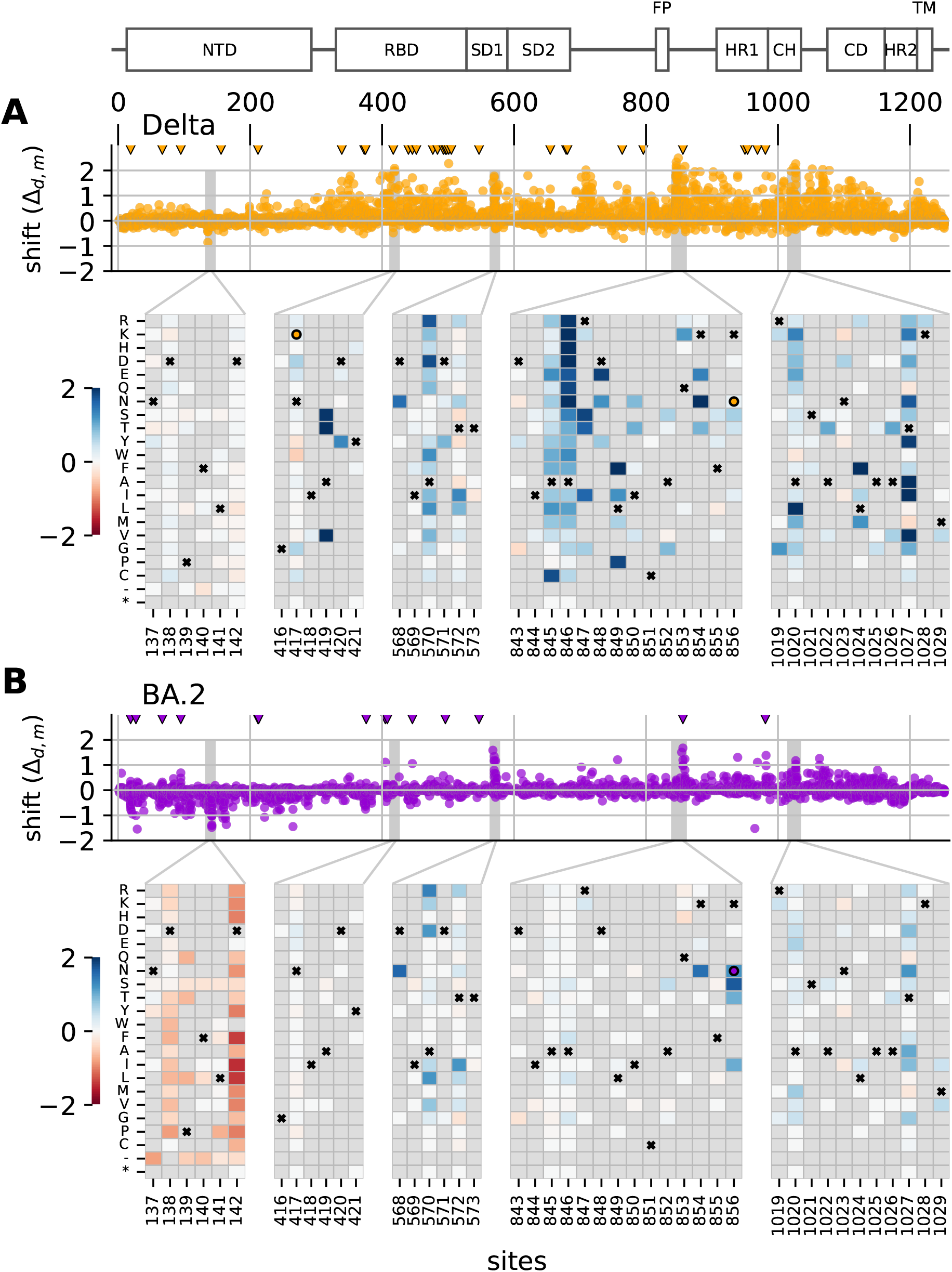
Distribution of shifts in spike’s primary sequence. Panels **(A)** and **(B)** show shifts for Delta and BA.2, respectively, relative to BA.1. The scatter plots show the values of all shift parameters at each site across spike’s primary sequence, with each dot corresponding to a parameter for a single mutation. Triangles at top mark the location of sites that differ in amino-acid identity in the homolog (Delta or BA.2) relative to BA.1. The diagram above the Delta’s scatter plot shows spike’s domain architecture. Heat maps show the shift parameters for individual mutations in key regions of spike with large shifts, with the color scale truncated at lower and upper limits of -2 and 2. Boxes with an “x” in the heat maps indicate the BA.1 amino-acid identity at a site. If the Delta or BA.2 wildtype amino acid differs from the BA.1 wildtype amino acid, then boxes with a circle indicate the Delta or BA.2 identity. A grey box indicates that the mutation was not observed in at least one of the three homolog DMSs; this is the case for many mutations as the libraries were largely designed only to include mutations observed among sequenced SARS-CoV-2 sequences (39).

### Experimental validation of mutational shifts

We experimentally validated the inferred shifts in mutational effects using spike-pseudotyped lentiviral particles (40). Specifically, we generated luciferase-expressing lentivirus pseudotyped with spikes carrying individual mutations inferred to have large shifts in mutational effects in the above analysis. We then measured the viral titer of these spike-pseudotyped lentiviruses in ACE2-expressing 293T cells and compared the titers of each mutant to the titers of an unmutated spike for each homolog, performing each experiment in triplicate (Figure 4).

**Fig. 4.**
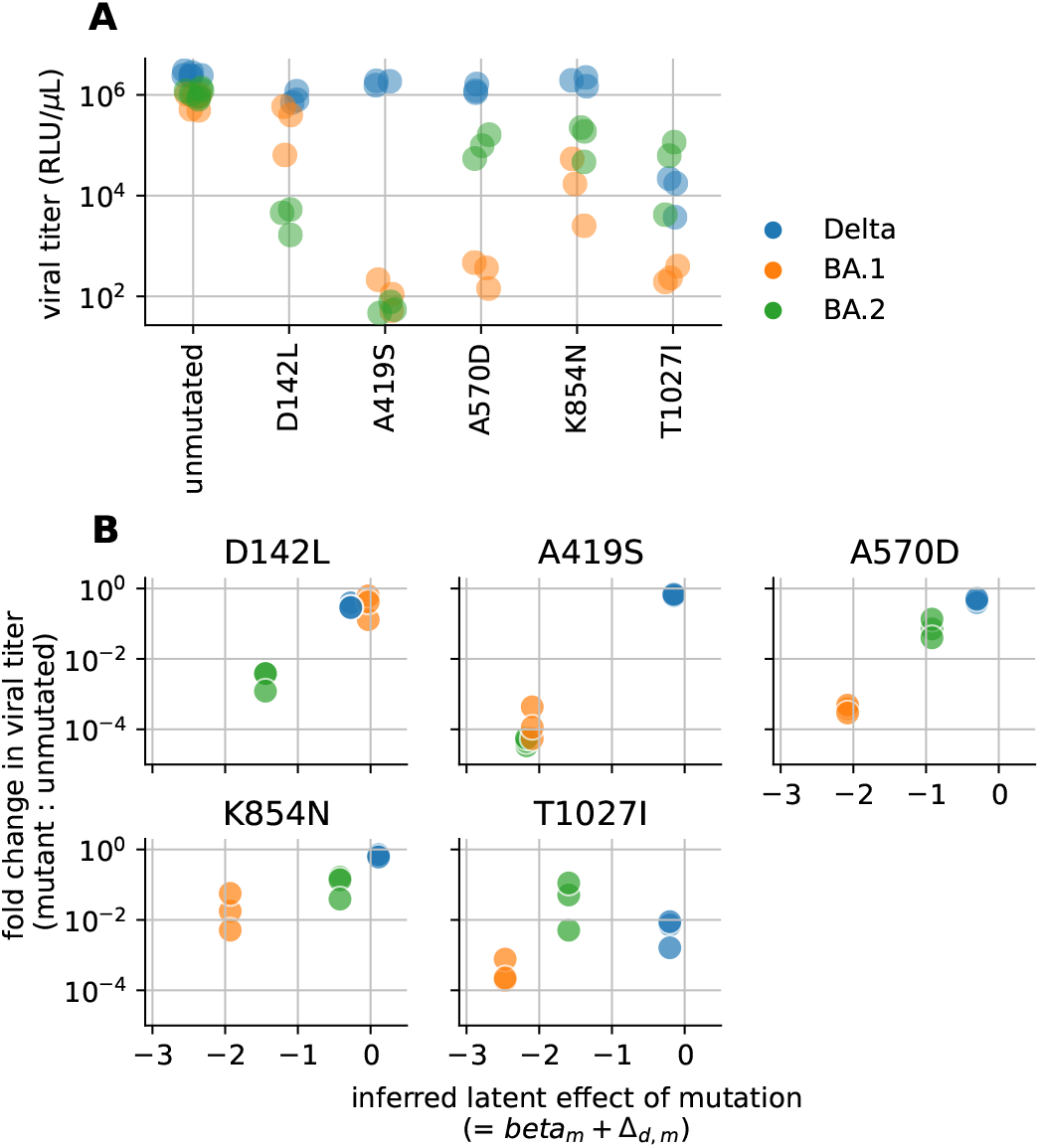
Experimental validation that a set of mutations inferred to have shifted effects indeed have different impacts on spike-mediated viral infection. **(A)** Titers of lentiviruses pseudotyped with the given spike variant, in relative light units (RLU) per *μ*L of virus. The points represent at least three independent replicates for each variant, and are colored by the spike homolog (Delta, BA.1, or BA.2). **(B)** Correlation of the predicted effect of each mutation in each spike homolog versus the actual experimentally measured impact of that mutation on spike-mediated viral infection. Each panel shows data for a different mutation. The y-axis shows the fold change in viral titer (from panel A) caused by the mutation relative to the unmutated spike homolog. The x-axis shows the inferred latent mutational effect of each mutation in each genetic background, expressed as the mutation’s effect in the reference background (*β*_*m*_) plus the mutation’s shift (∆_*d,m*_).

As expected, some mutations inferred to have undergone large shifts in their effects caused large changes in viral titer in some homologs (Figure 4A). In general, the changes in viral titers for different homologs were well correlated with the inferred shifts from the joint modeling of the DMS data (Figure 4B).

The most striking shift was for mutation A419S, which is highly deleterious in BA.1 and BA.2 (causing a *>*1,000-fold drop in titer), but is nearly neutral in Delta. The mechanistic basis of this shift is easily understood: A419S introduces an Nlinked glycan at 417 in BA.1 and BA.2 (which have N417), but not in Delta (which has K417). This glycan greatly reduces ACE2 affinity (16, 17), making A419S highly deleterious in BA.1 and BA.2 but not Delta.

The other validated mutations showed a similar pattern, where each mutation was highly deleterious in at least one homolog and substantially less deleterious in at least one other, with specific patterns differing by mutation. In nearly all cases, the inferred shifts in latent phenotype matched the experimentally measured shifts in effects on viral titer. These mutations occur in multiple regions of spike. D142L is in one of multiple loops in the NTD that form an antigenic supersite (41) and help modulate the efficiency of spike-mediated cell entry (42). The A570D and K854N mutations are both within a region in spike’s structure that regulates the balance between the up and down conformations of the RBD (Figure 5A) (43). The A570D mutation was proposed to be a key mutation that altered this up/down balance in the Alpha variant (43). The T1027I mutation is in the central helix, which forms part of spike’s trimerization interface. The mechanistic basis of these other validated shifts is less clear to us. But, together, they suggest that there have been large shifts in mutational effects relating to multiple functional and structural properties of spike.

**Fig. 5.**
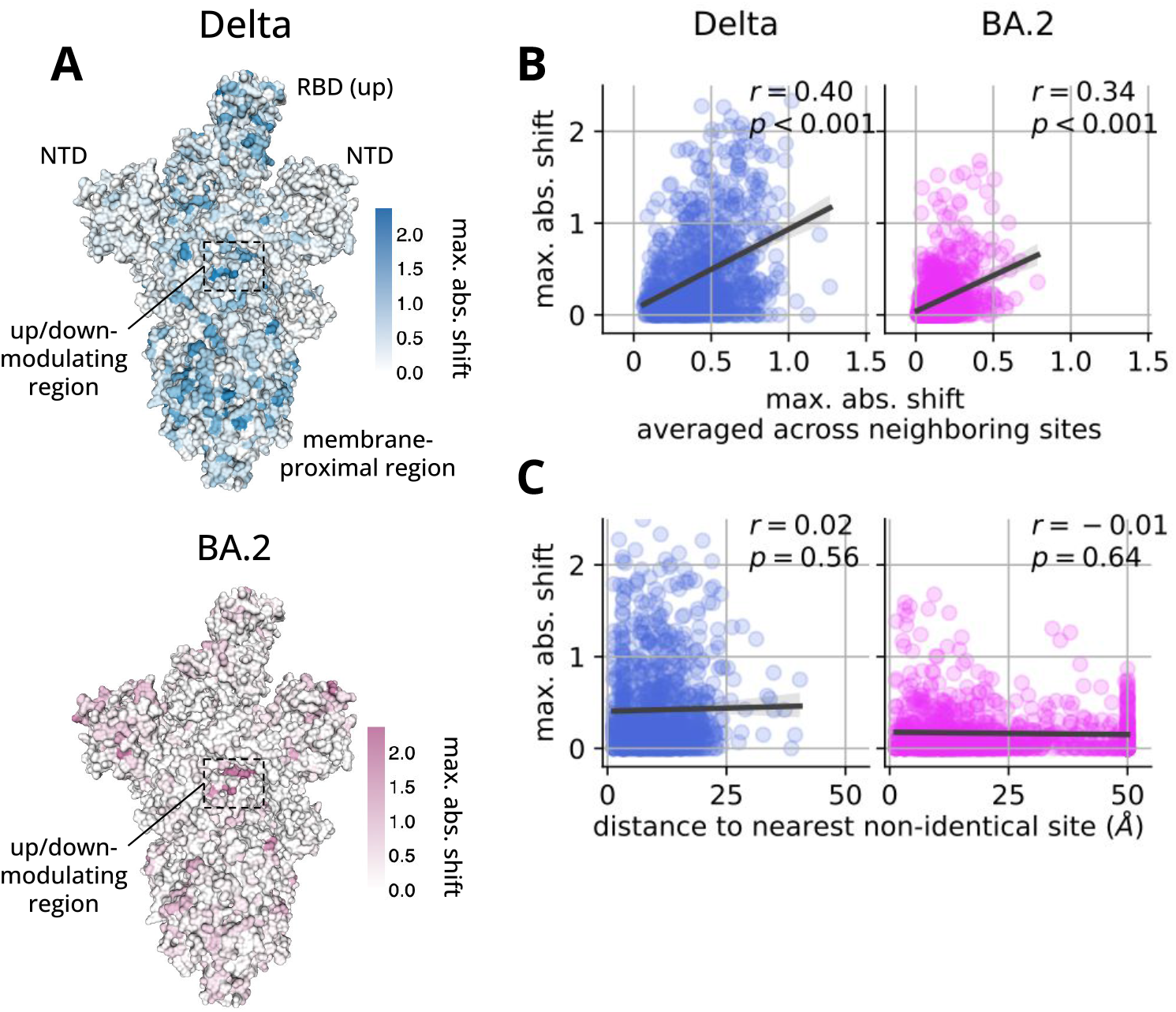
Distribution of shifts in spike’s threedimensional structure. **(A)** The trimeric ectodomain of spike with a single RBD in the up conformation (PDB 7TL9 (46)). The surface of the structure is colored by the maximum absolute value of all shift parameters at a given site, with the top and bottom images showing data for Delta and BA.2, respectively. Text indicates the location of the two NTDs and one RBD that are readily visible from this angle, as well as the up/down-modulating region that includes sites 568-573 and 843-856 from Figure 3. Images created using dms-viz (https://github.com/dms-viz). **(B)** We analyzed data across three different structures of spike with either zero, one, or two RBDs in the up conformation (PDB 7TF8, 7TL9, 7TGE (46)). For each pair of sites in spike’s primary sequence, we computed the minimum distance between those sites in the above structures, considering all heavy atoms from the corresponding residues. In the plots, each dot corresponds to an individual site, where the yaxis shows the maximum absolute value of all shift parameters at a given site and the x-axis shows this value averaged across all neighboring sites, where sites are considered neighbors if the minimum distance between them is less than 5Å. The left and right plots show data for Delta and BA.2, respectively, and *r* and *p* report the Pearson correlation coefficient and corresponding p-value. In each plot, the data are positively correlated, indicating that sites with large shifts tend to occur near other sites with large shifts. **(C)** Same as panel B, but the x-axis now shows the minimum distance of a given site to the nearest non-identical site between BA.1 and the given homolog, clipped at a maximum value of 50Å.

The validated mutations were often deleterious in BA.1 and considerably less deleterious in Delta due to large positive shifts. In general, shifts in Delta tended to be positive (Figure 3), and the corresponding shifted mutations tended to be deleterious in BA.1 (Figure S7), suggesting that Delta might be more mutationally tolerant than BA.1. Indeed, BA.1’s spike, and the monomeric version of its RBD, were found to be substantially less stable than those from the original D614G strain (44), which could lead to a lower tolerance for mutations (32). However, at least part of this bias could come from experimental artifacts (e.g., purifying selection was weaker in Delta’s DMS experiments than BA.1’s and BA.2’s; Figure S2).

In an effort to further validate the inferred shifts, we compared our inferences to ones that we computed from other studies, including DMS experiments of RBD homologs measuring mutational effects on ACE2 binding and RBD expression on the surface of yeast (16, 17), and a computational study that estimated mutational effects by analyzing millions of sequenced SARS-CoV-2 genomes from nature (45). Mutational effects in BA.1 and shifts in Delta and BA.2 relative to BA.1 were correlated between our study and these studies (Figure S8), lending additional support to our inferences. The other studies also suggest that Delta might be more mutationally tolerant than BA.1 (Figure S7).

### Distribution of shifts in spike’s structure

Sites with strongly shifted mutations tended to cluster in three-dimensional space (Figure 5A and B). We hypothesized that this clustering was occurring near sites that are non-identical between homologs, since differences in wildtype amino acids could lead to shifts at neighboring sites from short-range epistatic interactions. Surprisingly, although some sites with large shifts were structurally adjacent to non-identical sites, many were not (Figure 5C), suggesting that many shifts are due to long-range epistatic interactions between a shifted site and one or more non-identical sites. Thus, during spike’s evolution, mutations to one part of the protein can change its mutational tolerance elsewhere in the protein in unpredictable ways. A similar trend was seen in a study of shifts in mutational effects between homologs of HIV’s envelope protein (14). Both SARS-CoV-2 spike and HIV envelope are highly complex and conformationally dynamic proteins, which may facilitate such long-range interactions.

## Discussion

We describe a general method for jointly fitting a single model to multiple DMS experiments to identify mutations that have shifted effects across homologs or selective conditions. Algorithmically, the method is essentially an extension of globalepistasis models (31) to multiple experiments, which is useful because it allows the model to directly assess whether apparent shifts are strongly supported by the data from each experiment. We show the method can be used to identify shifts in mutational effects among three homologs of SARS-CoV-2 spike. The inferences in the model validate extremely well in experiments, with some mutations having effects on spike-mediated viral infection that differ by *>*1,000 fold between homologs. We also demonstrate that the shifts inferred using the joint-modeling approach are more consistent between replicates than ones inferred by separately modeling each experiment, suggesting that the joint-modeling approach is more effective at extracting real biological signal from noisy experiments.

Our method does make several assumptions. First, the joint-modeling approach assumes that most mutations have similar effects between experiments. This approach would not make sense if many mutations are expected to have large shifts, which can occur when comparing highly divergent homologs (18, 19). Second, by modeling all experiments on the same global-epistasis curve, the approach assumes that functional scores are directly comparable between experiments. This is not guaranteed. For instance, enrichment ratios are usually computed relative to the wildtype sequence from a given experiment. If the experiments have different wildtype sequences, the resulting enrichment ratios will systematically differ between experiments, which poses a problem if the wildtype sequences have large fitness differences. This did not appear to be a significant issue when comparing the spike homologs in this paper, perhaps because each wildtype spike homolog is roughly equally proficient at supporting the entry of pseudoviruses into cells (Figure 4A). However, in future use cases where this might be an issue, the *SI Appendix* and online documentation for `multidms` suggest strategies for normalizing functional scores between experiments to help make them comparable.

Despite the above assumptions, we envision that our method could be applied to many future studies comparing DMS experiments. Most DMS experiments have appreciable levels of noise, necessitating a method to account for noise when comparing them. Our method enables the use of globalepistasis models to analyze libraries with multiple mutations per variant, but is also compatible with libraries that only have a single mutation per variant (see *SI Appendix*). Further, we developed an open-source software package with comprehensive documentation so that others can easily use our method. Altogether, this method could greatly accelerate future use of DMS to identify mutations with biologically interesting shifts in effects.

## Materials and Methods

### Data and code availability and reproducibility

The `multidms` Python package is available via the Python Package Index (PyPI). It provides tools for processing functional scores from DMS data, fitting a `multidms` model to the data, and generating plots for analyzing the results. The core models and optimization algorithms are implemented using the JAX and JAXopt packages, enabling automatic differentiation and just-in-time compilation for high performance on CPU and GPU (47, 48). The source code is available and maintained at https://github.com/matsengrp/multidms, and carries an MIT license. For further details on installation, interface, how to contribute and more, see our package documentation (https://matsengrp.github.io/multidms/).

For the full analysis pipelines used to generate functional scores from raw DMS data for a given spike homolog, see:

1. Delta (39): https://dms-vep.github.io/SARS-CoV-2_Delta_spike_DMS_REGN10933/
2. BA.1 (39): https://dms-vep.github.io/SARS-CoV-2_Omicron_BA.1_spike_DMS_mAbs/
3. BA.2 (from this study): https://dms-vep.github.io/SARS-CoV-2_Omicron_BA.2_spike_DMS/

We created a GitHub repository (https://github.com/matsengrp/SARS-CoV-2_spike_multidms) with all code used to curate the above DMS data, fit `multidms` models to these data, and make all figures in the paper. The code, as well as a step-by-step explanation of the analysis pipeline, is in a single Jupyter Notebook, an HTML version of which can be viewed at https://matsengrp.github.io/SARS-CoV-2_spike_multidms/ (49). The repository also includes all input data, key ouptut files, and instructions for running the notebook. Finally, we make the shift parameters (∆_*d,m*_ accessible at https://matsengrp.github.io/SARS-CoV-2_spike_multidms/ spike-analysis.html#shifted-mutations-interactive-altair-chart as an interactive version of Figure 3 made using Altair (50).

### Jointly modeling multiple DMS experiments

We now define notation and introduce our model more formally. Let *M ∈* ℕ denote the number of distinct mutations, and represent a given variant *v ⊂ M≡ {*1, …, *M}* as an index set of the mutations it contains (*v* is in the set of subsets of the *M* mutations, i.e. *v ∈ 𝒱≡* 2^*ℳ*^, where 2^*ℳ*^ denotes the power set of ℳ). We depart from the informal main text notation and represent a variant *v* as an indicator (one-hot) vector *x*_*v*_ *∈* {0, 1} ^*M*^ where [*x*_*v*_]_i_ = 1 if *i ∈ v* and [*x*_*v*_]_*i*_ = 0 otherwise. We will express the model in vector/matrix notation, rather than the element-wise notation used in the main text model summary (we use column vectors by convention).

Let *D ∈* ℕ be the number of experiments (the letter *D* is used as a mnemonic for DMS) and write 1_*D*_ for the *D*-vector of ones. We introduce an additive *latent phenotype* model jointly for *D* experiments via a family of affine maps 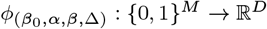 defined by

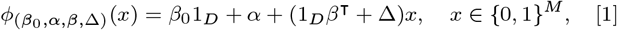

where the family is parameterized by intercept *β*_0_ ∈ ℝ and mutational effects *β ∈* ℝ^*M*^ that are shared by all *D* output dimensions, global offset *α ∈* ℝ^*D*^, and shift matrix ∆ ∈ ℝ^*D*×*M*^. We require that the first row of ∆ is the zero *M* -vector and the first element of *α* is zero, so that the reference experiment (indexed 1 WLOG) has no shifts, and *β* is then interpreted as the vector of mutational effects in the reference experiment, with the intercept *β*_0_ representing the latent phenotype of the wildtype sequence in the reference experiment.

Next, we introduce a *global-epistasis function* via a family of strictly monotone maps *g*_*θ*_ : ℝ → ℝ that we use to take latent phenotypes to predicted functional scores. This family is parameterized by *θ ∈* ℝ^*r*^ for some *r ∈* ℕ. For the results presented in this study,we use the sigmoid function

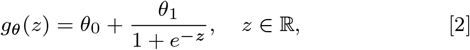

with *r* = 2 parameters, which allows us to adapt the output range of the global-epistasis function (the interval (*θ*_0_, *θ*_0_ + *θ*_1_)) to the range of our functional score data, but is otherwise a fixed link function (imposing a *gauge* on our latent phenotype model parameters). We finally compute the predicted functional score in experiment *d ∈ {*1, …, *D}* of a variant *v ∈ V* with one-hot encoding *x*_*v*_ *∈ {*0, 1*}*^*M*^

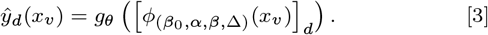

### Inferring model parameters from the DMS data

Our data consist of sets of one-hot encoded variants and their associated functional scores from each of *D* experiments. Denote these as *D*_*d*_ *⊂ {*0, 1*}*^*M*^ *×* ℝ for *d* = 1, …, *D*. We minimize an objective of the form

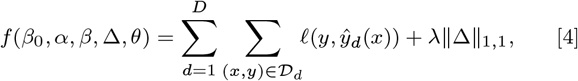

where *𝓁* : ℝ × ℝ → ℝ is a Huber loss function measuring the difference between a predicted and an observed functional score, *λ ∈* ℝ is the lasso penalty weight, and ∥ · ∥_1,1_ denotes the entrywise *L*_1_ norm (not to be confused with the matrix 1-norm ∥ · ∥_1_). Note that the parameters (*β*_0_, *α, β*, ∆, *θ*) appear in the loss function via Eq. (3), but are suppressed in Eq. (4) for notational compactness. Note also that by taking *λ* = 0, the loss term of the objective becomes separable over the *D* experiments, so marginal inference is recovered as a special case.

For a general global epistasis function *g*_*θ*_, the objective Eq. (4) is in general non-convex. However, with the simple sigmoid function Eq. (2), it is bi-convex in (*β*_0_, *α, β*, ∆) and *θ*. This can be seen by noting that, for fixed *θ*, the prediction model takes the form of a generalized linear model with a sigmoid link function, and for fixed (*β*_0_, *α, β*, ∆), the model parameterized by *θ* is a linear regression problem. The objective Eq. (4) has a smooth loss term and a non-smooth penalty term. We minimize it using the Nesterov-accelerated proximal gradient method with backtracking line search (51), taking gradient steps using the smooth term, and applying a proximity operator associated with the non-smooth term.

### Deep mutational scanning of SARS-CoV-2 spike

Delta and BA.1 full spike DMS libraries were designed as described previously in (39). BA.2 full spike deep mutational scanning libraries were designed using the same methods as BA.1 libraries except using BA.2 spike as a template sequence. The sequence of BA.2 spike can be found at https://github.com/dms-vep/SARS-CoV-2_Omicron_BA.2_spike_DMS/blob/main/library_design/reference_sequences/3332_pH2rU3_ForInd_Omicron_sinobiological_BA2_B11529_Spiked21_T7_CMV_ZsGT2APurR.gb. In each library, a random 16-nucleotide barcode was included downstream of the stop codon of each spike variant, such that each variant is associated with a unique barcode. Long-read PacBio sequencing was used to acquire reads spanning the entire spike gene and barcode, allowing a variant’s genotype to be associated with its barcode, as described in (39). Many spike variants appeared multiple times in a given library, associated with multiple unique barcodes.

DMS functional selections were performed as described previously in (39). In brief, 1 million HEK-293T-ACE2 cells were infected with 0.6-1 million spike-pseudotyped library variants or 5 million of VSV-G pseudotyped variants. 12-15 hours post infection, cells were trypsinized, washed with PBS, and non-integrated viral DNA was extracted using QIAprep Spin Miniprep Kit. Extracted DNA was used to prepare PCR amplicon libraries for Illumina sequencing. Libraries were sequenced using NextSeq 2000 P2 and P3 reagent kits. The resulting data provided counts for each 16-nucleotide barcode in each sample.

For each experiment, a functional score was computed for each barcoded variant as a log-enrichment ratio: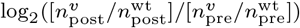, where 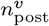gives the number of deep-sequencing counts for variant *v* in the post-selection library (from cells infected with spike-pseudotyped viruses), 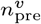 gives counts for *v* in the pre-selection library (from cells infected with VSG-G-pseudotyped viruses), and 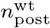 and 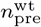 give these same counts but for the wildtype homolog from a given experiment. A pseudocount of 0.5 was added to each of these counts to avoid dividing by zero. Negative functional scores indicate that a given variant was depleted relative to wildtype, while positive functional scores indicate that the variant was enriched relative to wildtype.

### Fitting a multidms model to the spike DMS data

To start, we curated functional scores described in the above section in the following ways. To reduce noise, we discarded data for all barcoded variants with fewer than 100 pre-selection counts. Due to experimental batch effects, the range of functional scores differed between experiments (data not shown). For instance, variants with stop codons tended to have more negative functional scores in the BA.1 and BA.2 experiments compared with Delta. To help make scores more comparable between homologs, we truncated all functional scores from all experiments at a lower bound of -3.5 and an upper bound of 2.5 (Figure S2B). The lower bound of -3.5 roughly corresponds to the lower end of the dynamic range of the assay. Although functional scores can go below this number, how negative a functional score can get is partially determined by experiment-specific factors such as deep-sequencing depth.

The DMS experiments were performed with at least two biological replicates per homolog, where each replicate experiment used an independently synthesized barcoded variant library. Each of the Delta and BA.2 replicate experiments were performed with two technical replicates, and we combined all functional scores between pairs of technical replicates into a single dataset. For variants associated with multiple unique barcodes in a single biological replicate dataset, we averaged the variant’s score across all barcodes. This averaging step increased the speed of model fitting without substantially changing the final results (data not shown).

Some sites in the spike protein were mutated in one or two of the homolog DMS libraries, but not all three. For instance, due to indels, some sites that are present in one homolog are completely missing in another. Since it is not possible to compute shifts across all homologs at such sites, `multidms` automatically discards all variants with mutations at any of these sites.

We fit a single `multidms` model to one biological replicate dataset per homolog, using BA.1 as the reference, and using 30,000 proximal gradient iterations to allow the Huber loss term to converge (Figure S9). We then independently fit a second model to a second set of biological replicate datasets. Figure S10A shows the sigmoidal global-epistasis function inferred in each replicate fit at a regularization weight of *λ* = 5 ×10^*−*5^ (the next section describes our logic for choosing this weight). Most data fit to the lower end of the sigmoid, suggesting the model is capturing saturating effects of deleterious mutations. Observed functional scores from the training data were well correlated with predicted scores for each experiment from each replicate (Figure S10B).

### Choosing a regularization weight

The *Results* section reports data from `multidms` models fit using a regularization weight of *λ* = 5 ×10^*−*5^. Below, we describe our strategy for choosing this weight. We tested several weights that ranged between zero and 0.001, fitting one model per weight. As expected, increasing the weight tended to shrink the inferred shift parameters, with some parameters shrinking more rapidly than others. Figure S4A shows examples of this pattern for different sets of mutations. The red lines show patterns for mutations to stop codons. The effects of these mutations are not expected to be shifted between homologs as they should be equally deleterious in each. At very small weights, some stop mutations were inferred to have large non-zero shifts, presumably due to experimental noise in the data. However, as the weight is increased, these shifts are driven to zero, with nearly all shifts reaching zero by the time the lasso weight reaches *λ* = 5 ×10^*−*5^ (Figure S4B). In contrast, there are some shifts that are not driven to zero for this value of *λ*. For example, the five nonsynonymous mutations that we experimentally validated to have large shifts are only driven to zero by much larger weights (see colored lines). Such shrinkage patterns of the validated mutations were highly consistent between replicates.

We also compared weights based on the model’s ability to predict experimentally measured functional scores in the training data, as quantified by the loss function used to train the model, not including the lasso term (Figure S4C). As expected, the loss increased as the lasso weight increased. At lower weights, this increase was gradual, before becoming steeper at intermediate weights and leveling out at the highest weights. The steepest increases came for *λ* > 5 ×10^*−*5^. Together, the above results show that a lasso weight of *λ* = 5 ×10^*−*5^ was needed to drive shifts for stop codon mutations—a rough proxy for noise—to zero, but that higher weights resulted in substantially worse loss, suggesting over-regularization.

We also quantified the correlation of shift parameters between the replicate model fits as a function of lasso weight (Figure S4D). In each fit, the model from one replicate has never seen the data used to fit the model from the second replicate, and vice versa. The correlation in shift parameters tends to increase as *λ* is increased from 0 to 5 ×10^*−*5^. This pattern is consistent with the hypothesis that the shift parameters from each replicate are overfit to their corresponding datasets at low weights, and that increasing the weight tends to reduce overfitting, leading to a higher correlation. Of note, at the highest tested weights, mutations that we experimentally validated to have large shifts were inferred to have shifts near zero, indicating that these weights are too strong. The correlation of *β*_*m*_ parameters between replicate fits remained high across all *λ* values, showing that increases in *λ* can dramatically improve the correlation for shift parameters while retaining a high correlation for *β*_*m*_ parameters (Figure S4D).

In all, the above lines of evidence suggest that a lasso penalty *λ* = 5 ×10^*−*5^ was sufficient to suppress noise, while preserving biologically relevant signal.

### Experimental validation

Spike genes with desired mutations were introduced using PCR with overlapping mutation-carrying primers followed by HiFi assembly. Plasmids used as Delta, BA.1 and BA.2 spike templates can be found at https://github.com/dms-vep/SARS-CoV-2_Omicron_BA.2_spike_DMS/tree/main/library_design/plasmid_maps. Pseudoviruses were generated using a method described previously (40) with the following changes: pHAGE6_Luciferase_IRES_ZsGreen was used as the backbone for which only Gag/Pol helper plasmid and the spike expression plasmid are required to generate a virus. Produced pseudoviruses were titrated on HEK-293T-ACE2 by performing duplicate serial dilutions and virus titers were measured 48 hours after infection using Bright-Glo Luciferase Assay System (Promega, E2610).

## Acknowledgments

We thank Gabriel Boyle and Daniel Ellis for useful discussions. This work was supported in part by the NIH under the grants R01 AI162611 and R01 AI141707, and the Genomics & Bioinformatics Shared Resource, RRID:SCR_022606, of the Fred Hutch/University of Washington Cancer Consortium (P30 CA015704). F.A.M and J.D.B. are investigators of the Howard Hughes Medical Institute. W.S.D. was supported by a Fellowship in Understanding Dynamic and Multi-scale Systems from the James

S. McDonnell Foundation. Scientific Computing Infrastructure at Fred Hutch funded by ORIP grant S10OD028685.

