## Supporting Information for "Jointly modeling deep mutational scans identifies shifted mutational effects among SARS-CoV-2 spike homologs"

### Supporting Information Text

#### Additional modeling details and options

Our model is summarized in the main text Results and Methods. Here, we provide additional details and modeling options that have been implemented in the `multidms` software package.

**Reckoning mutations with respect to the reference experiment.** If a non-reference experiment has a different wildtype sequence from the reference (i.e., the wildtype sequences are different homologs), the set of mutations at non-identical sites that convert between the two sequences are considered when computing latent phenotypes via Eq. (1) in the main text. We refer to this set of mutations as "the bundle". The bundle of mutations is also considered when computing latent phenotypes of variants from non-reference experiments.

If a mutation occurs at a non-identical site, then the mutation is always defined relative to the reference. For instance, say site 30 is a Y in the non-reference experiment's wildtype sequence and an A in the reference experiment's wildtype sequence. If a variant from the non-reference experiment had a Y30G mutation, then the mutation would be defined as A30G in the summation term. This does not assume that Y30G has the same effect as A30G, it is simply being consistent with a model that computes the latent phenotypes of all variants relative to a common reference sequence. This strategy ensures that each experiment informs the same set of mutation effects $\beta$ and shift parameters $\Delta$. If a variant from a non-reference experiment had a reversion mutation at a non-identical site (e.g., Y30A), then the mutation would not be included in the summation term of Eq. (1) for that variant since the variant would have the reference amino-acid identity at that site. In comparison, A30Y would be included for all variants from the non-reference experiment that lack a mutation at site 30.

If one or more of the mutations in the bundle are not included in the DMS libraries (e.g., Y30A is missing from the non-reference experiment and A30Y is missing from the reference experiment), then the individual $\beta_m$ and $\Delta_{d_m}$ parameters for these unmeasured mutations cannot be inferred from the data. To address this problem, we remove these parameters from the summation term of Eq. (1) in the main text, and use a single $\alpha_d$ parameter to infer the combined effects of these mutations. In this scenario, the use of $\alpha_d$ is important for allowing the model to learn the full latent phenotypic distance between wildtype sequences from different experiments. Otherwise, if all such mutations are sampled, $\alpha_d$ can be locked at zero.

If there is an unmeasured mutation in the bundle, our strategy for removing this mutation from the summation term of Eq. (1) is to completely ignore this site when analyzing the data. This involves ignoring all variants with mutations at this site. Although this can result in throwing away data, it is necessary for the math to work out. For instance, in the above example where Y30A is missing from the non-reference experiment and A30Y is missing from the reference experiment, $\alpha_d$ will be used to capture the effect of A30Y when modeling the latent phenotype of the wildtype sequence from the non-reference experiment. If a variant from the non-reference experiment has a Y30G mutation, this mutation will be included in the summation term of Eq. (1) as A30G. However, since $\alpha_d$ is a constant offset used to compute the latent phenotype of *all* variants from experiment $d$, and since $\alpha_d$ includes the effect of A30Y, Eq. (1) would effectively be adding the effects of both A30G and A30Y when computing the latent phenotype of this particular variant, which does not make sense. Because of this problem, we choose to ignore all data at such sites.

**Parameterizing global epistasis.** The `multidms` software allows users to choose various functional forms for the nonlinearity $g$, in addition to the sigmoidal function used in the main text Eq. (2).

- Sigmoid:

$$g(z) = \frac{\theta_{\text{scale}}}{1 + e^{-z}} + \theta_{\text{bias}}$$

  where $\theta_{\text{scale}}$ is a scaling factor that allows the sigmoid to stretch or contract along the y-axis and $\theta_{\text{bias}}$ is a factor that allows it to translate along the y-axis. We do not include analogous factors for the x-axis, since these factors would be redundant with the affine transformation parameters in Eq. (1). This choice can be used to model a Hill equation with a Hill coefficient of 1.

- Softplus:

$$g(z) = -\theta_{\text{scale}} \log\left(1 + e^{-z}\right) + \theta_{\text{bias}}$$

  This function is a log-transformed version of a sigmoid function from above, with $\theta_{\text{scale}}$ and $\theta_{\text{bias}}$ parameters serving similar roles. This function may be more appropriate for modeling functional scores in log space.

- Single-layer neural network with sigmoid activations:

$$g(z) = b^{(o)} + \sum_{i}^{n} \frac{w_i^{(o)}}{1 + \exp\left(w_i^{(h)} z + b_i^{(h)}\right)}$$

  where $n$ is the number of units in the hidden layer and is defined by the user; $w^{(h)} \in \mathbb{R}^n_{\geq 0}$ and $b^{(h)} \in \mathbb{R}^n$ are the hidden unit weights and biases, respectively; and $w^{(o)} \in \mathbb{R}^n_{\geq 0}$ and $b^{(o)} \in \mathbb{R}$ are the (linear) output weights and bias, respectively. All weights in the network are clipped to a minimum of zero during training to ensure monotonicity in the resulting function. We caution that this is an advanced feature, and that choosing large $n$ may result in overfitting to outliers in the data.

- Identity:

$$g(z) = z$$

Here there is no global epistasis—latent phenotypes are identical to functional scores. We recommend using this if none of the variants in the DMS experiment have more than one mutation, since global-epistasis is a manifestly multi-mutant phenomenon. It can also be used as a baseline against which to compare non-linear global-epistasis functions.

**Functional score alignment.** Since the model uses a single global-epistasis model to jointly model all experiments, it assumes that the functional scores are directly comparable between experiments. In practice, this is not always the case. Below, we describe two optional features to help make the scores more comparable.

***Optional normalization of observed functional scores.*** A common way to compute functional scores is with log-enrichment ratios, where all scores from a given experiment are normalized so that the wildtype sequence from that experiment has a value of zero. If wildtype sequences differ between experiments, then log-enrichment ratios may not be directly comparable between experiments, as they are normalized to different reference points. This breaks the assumption of our joint-modeling scheme: that all functional scores are directly comparable.

Ideally one or more of the same sequences would be included in the DMS library of each experiment, so that all scores could be normalized to the same sequence. However, if there are no common sequences, then it may be possible to computationally estimate how to normalize scores so that they are more directly comparable.

To this end, the model includes an additional parameter $\gamma\mathbb{R}^D$ that allows functional scores from each experiment to be normalized as follows:

$$y_{v,d}^{\text{norm}} = y_{v,d} + \gamma_d$$

where $\gamma_d$ for the reference experiment is locked at zero. There is a theoretical basis for adding $\gamma_d$ to $y_{v,d}$ if functional scores are log-enrichment ratios. As mentioned above, log-enrichment ratios are normalized so that the wildtype sequence from a given experiment has a value of zero, according to the formula:

$$y_{v,d} = \log(E_{v,d}) - \log(E_{\text{wt},d})$$

Thus, adding $\gamma_d$ to $y_{v,d}$ is akin to renormalizing the log-enrichment ratios so that a different sequence has a functional score of zero. $\gamma$ can be used to recalibrate the functional scores of each non-reference experiment to the wildtype sequence of the reference experiment. If these values are not experimentally measured, the model allows $\gamma$ to be fit during optimization. Alternatively, $\gamma$ can be locked at zero if the user does not wish to implement this optional feature.

***Optional truncation of predicted functional scores.*** Experimentally measured functional scores might be constrained by the dynamic range of the assay. For instance, if the functional scores are log-enrichment ratios, the experiment may not be able to accurately measure scores below $-3$, in which case the user may truncate observed scores at this floor. When fitting the model to these data, it may be desirable to truncate *predicted* functional scores in a similar way. This is especially relevant when using nonzero $\gamma$ parameters. For instance, say the user has truncated observed functional scores at a lower bound of $-3$ across all experiments. If $\gamma_d$ is fit to $-1.0$ for a non-reference experiment $d$, then the new floor of normalized functional scores for that experiment would be $-4$, while the floor for the reference experiment would still be $-3$. In this case, the global-epistasis function becomes problematic: if it allowed predictions to go below $-3$, it could help model the floor of variants in the non-reference experiment, but hurt with modeling the floor of variants in the reference experiment. However, if it was able to truncate predicted scores for a given experiment at the floor for that experiment, then this tension would be relieved: predicted scores for the reference experiment could be truncated at $-3$, while (non-truncated) predicted scores for the non-reference experiment could go as low as $-4$.

To this end, the software package provides an option to truncate predicted functional scores. To enable this, the mathematical model passes predicted functional scores through an activation function, $t$. Users can choose between multiple options for $t$, one of which (*softplus*) truncates predicted functional scores at a user-specified lower bound, and the second of which (*identity*) leaves the functional scores unaltered. The functional forms of these options are as follows.

- Softplus:

$$t(y) = 0.1 \times \log\left(1 + e^{\frac{y-(l_d+\gamma_d)}{0.1}}\right) + l_d + \gamma_d$$

where $l_d$ is the floor of the dynamic range of the DMS experiment from experiment $d$, and is set by the user. Functionally speaking, this equation truncates scores at a floor of $l_d + \gamma_d$, while leaving scores above this floor (mostly) unaltered. There is a small range of input values where the function smoothly transitions between a linear regime (where data is not truncated) and a flat regime (where data is truncated). But, the 0.1 constants from above ensure a sharp transition between regimes. This function is differentiable and thus allows for gradient-based optimization.

- Identity:

$$t(y) = y$$

This function leaves functional scores unaltered, meaning scores are not truncated.

#### Supporting Figures

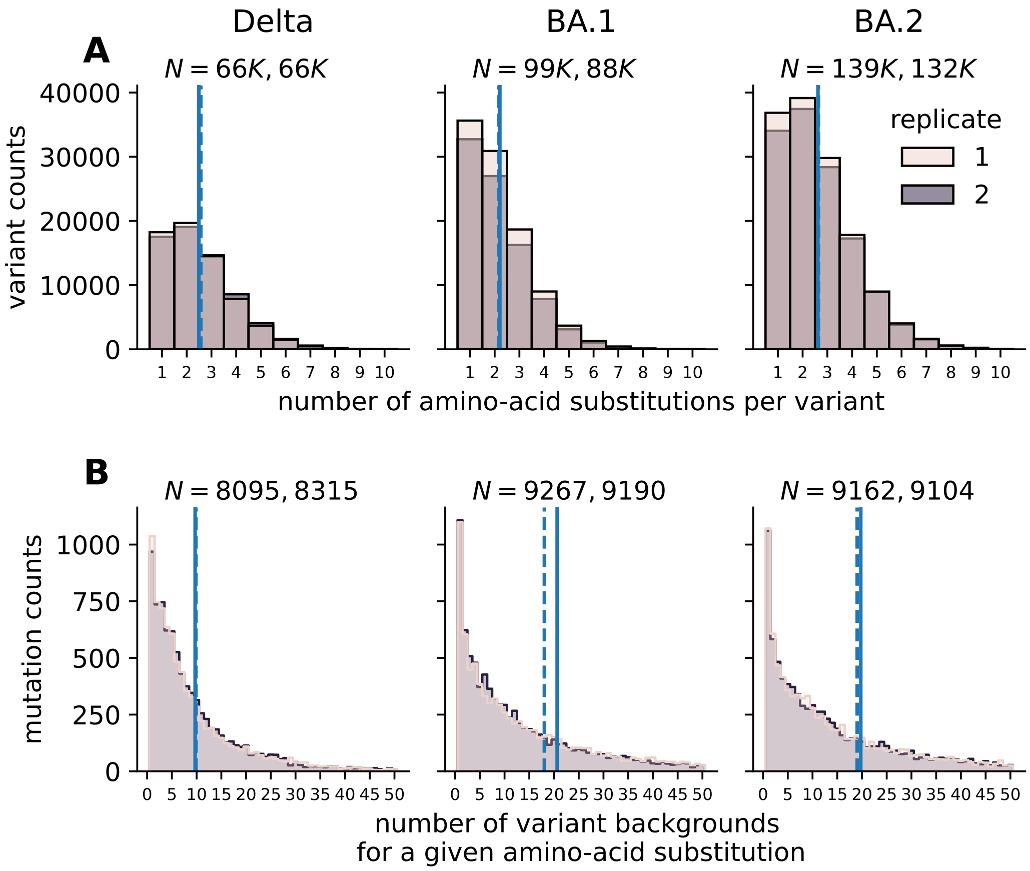


**Fig. S1.** Summary statistics of amino-acid substitutions and variants in unselected DMS libraries. Each distribution shows data for a given replicate DMS library (hue) for a given homolog (column). **(A)** Distribution of the number of amino-acid substitutions per variant, truncated at 10 counts. $N$ reports the number of variants in a given library, with one number for each of the two replicates. Blue lines show the mean of each distribution, with dashed and solid lines showing data for different replicates. **(B)** Distribution of the number of unique variants with a given amino-acid substitution, truncated at 50 counts.

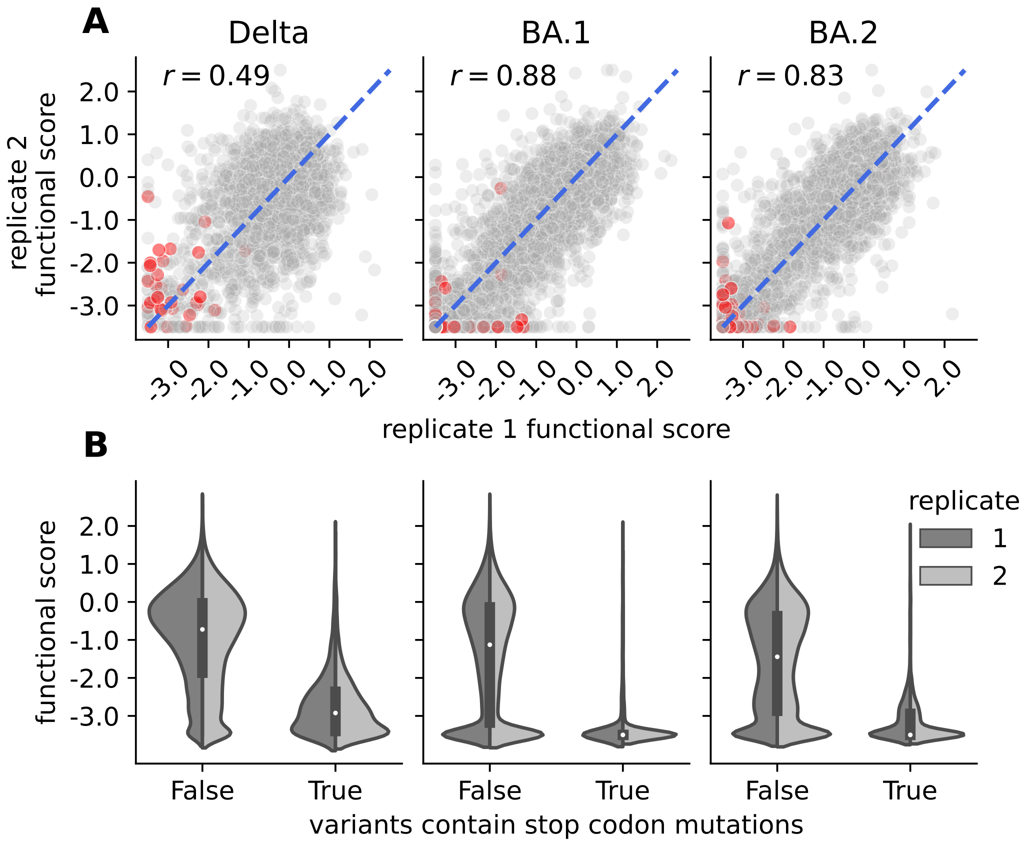


**Fig. S2.** **(A)** Correlation of functional scores between biological replicates of the DMS experiment performed on a given homolog. Each dot corresponds to a variant seen in both replicate experiments, which is a subset of all variants in a given experiment. Each plot reports the Pearson correlation coefficient. For each homolog, there was an intermediate correlation between replicates, suggesting that functional scores were reproducible, but also suggesting an intermediate level of noise, with Delta experiments being noisier than Omicron experiments. Red dots show data for variants with mutations to stop codons, while grey dots show data for all other variants. **(B)** Distribution of functional scores among variants with and without mutations to stop codons (x-axis) for a given replicate experiment (hue) performed on a given homolog (column). As expected, variants with stop codons tend to have lower functional scores than variants without stop codons. Notably, variants with stop codons tend to have somewhat higher functional scores in the experiments performed on Delta compared to BA.1 or BA.2, indicating that purifying selection may not have been as strong in the Delta experiments. This is an experimental artifact that may partially account for why inferred shifts tend to be positive in Delta relative to BA.1. In both panels, functional scores are clipped at a lower bound of -3.5, which we use as an approximation of the lower bound of the assay's dynamic range.

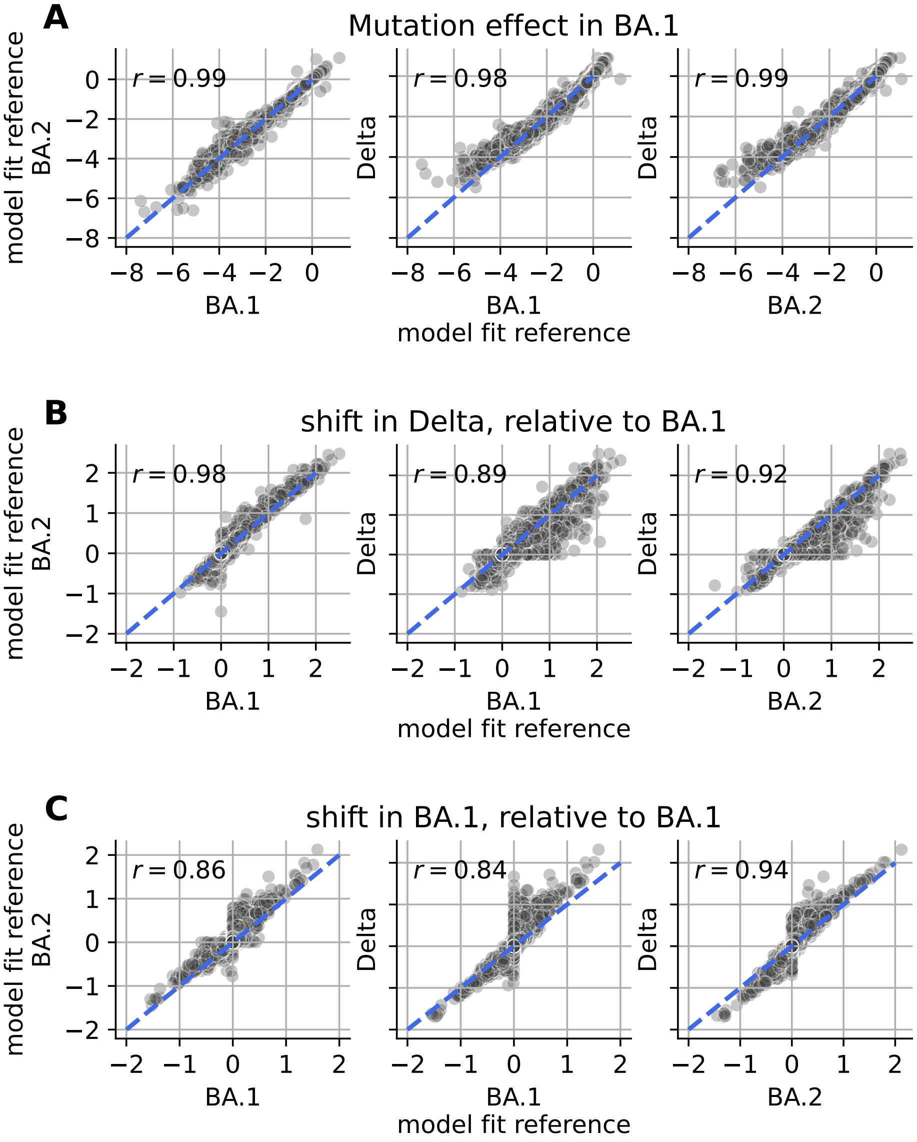


**Fig. S3.** Comparison of mutational effects and shift parameters inferred using different reference experiments. To generate these plots, we fit a `multidms` model to the experimental data as described in the main text, but using either the Delta, BA.1, or BA.2 experiment as the reference experiment. For each model, we computed the effect of each mutation in each homolog's experiment $d$ as $\beta_m + \Delta_{d,m}$, where $\Delta_{d,m}$ is always zero for the reference homolog from a given model. The following plots correlate results among models using a given homolog's experiment as the reference, always reporting shifts relative to mutational effects inferred in BA.1. $\mathbf{(A)}$ Correlates mutational effects in BA.1. $\mathbf{(B)}$ Correlates shifts in mutational effects in Delta relative to BA.1, computed by subtracting the effect inferred in Delta by the effect inferred in BA.1. $\mathbf{(C)}$ Correlates shifts in mutational effects in BA.2 relative to BA.1, computed by subtracting the effect inferred in BA.2 by the effect inferred in BA.1. $r$ reports the Pearson correlation coefficient and each dot corresponds to a different mutation. The effects inferred for BA.1 were highly correlated. The shifts in effects were also well correlated, but showed variability between models. Thus, the results were partially dependent on the choice of reference experiment, and suggest that future users of this method should also compare results between different choices. But the mutations with large shifts in one model almost always had large shifts in the other models, suggesting that the overall results were robust to this choice in the case of this study.

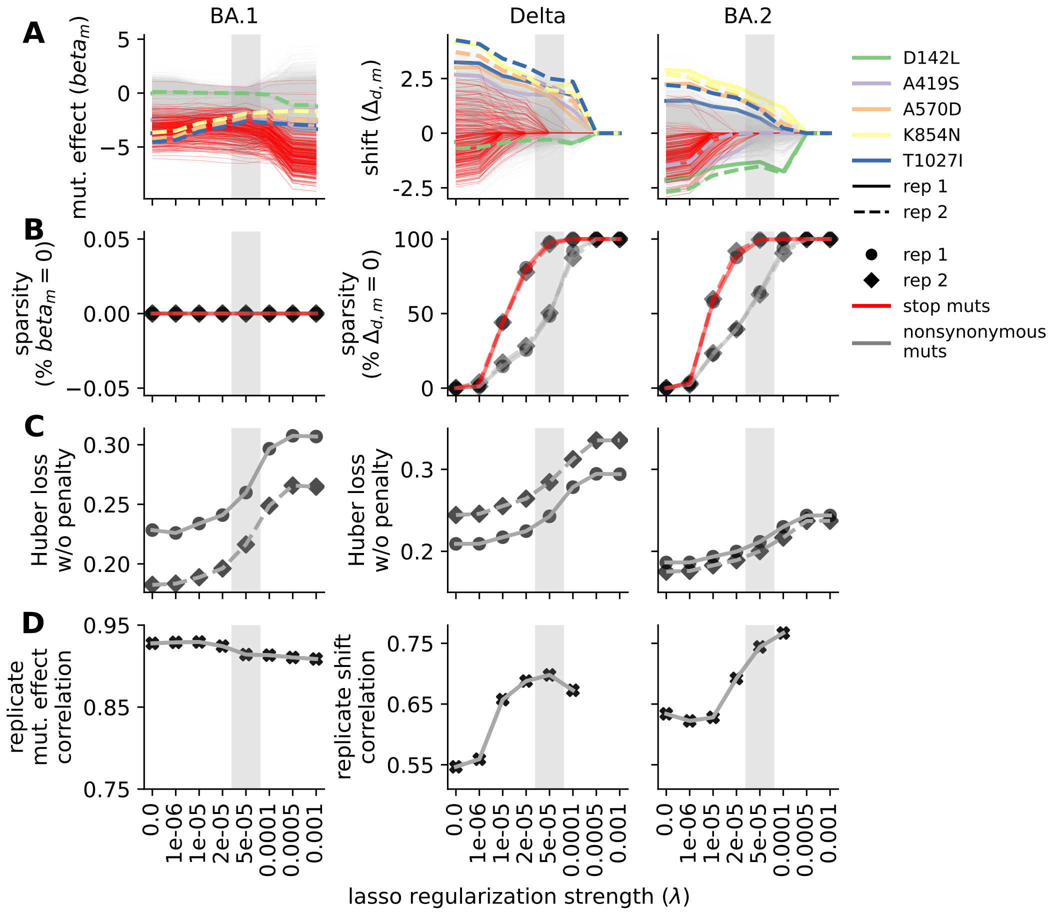


**Fig. S4.** To generate these plots, we fit a `multidms` model using each of the indicated lasso penalty strengths (x-axis). For each value, we fit two replicate `multidms` models on replicate input data, as described in the main text. **(A)** Lasso paths of $\beta_m$ parameters quantifying mutational effects in BA.1 (left column) or shift parameters quantifying shifts in mutational effects in Delta (middle column) or BA.2 (right column) relative to BA.1. Each line tracks the value of a $\beta_m$ or shift parameter (y-axis) for a single mutation as a function of lasso strength. Red lines show data for all mutations to stop codons. The other colored lines show data for the mutations specified in the legend, which includes all five mutations validated in Figure 4. Grey lines show data for all other mutations in the library. The dashed and solid lines show data for the two replicates. **(B)** The sparsity of each parameter set (i.e., the percent that equals zero) at a given lasso strength, with red lines showing data for mutations to stop codons and grey lines showing data for nonsynonymous and in-frame codon-deletion mutations. **(C)** The performance of the model at a given lasso strength, where performance is quantified as the Huber loss between predicted and experimentally observed functional scores in the training data for a given homolog (left plot: BA.1; center plot: Delta; right plot: BA.2). Lower loss corresponds to better performance. **(D)** Pearson's correlation coefficient of $\beta_m$ parameters (left plot) or shift parameters (center and right plots) from replicate model fits at a given lasso strength. For shift parameters, correlations are not shown for high weights where nearly all shift parameters are zero.

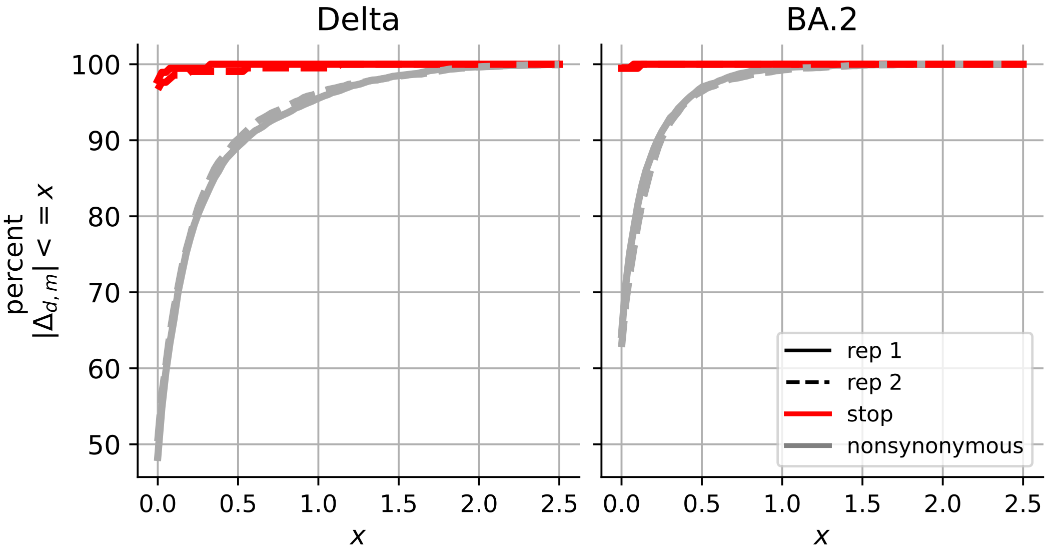


**Fig. S5.** Percent of shift parameters (y-axis) that are less than or equal to a given magnitude (x-axis). The red lines show data for mutations to stop codons, while the grey lines show data for nonsynonymous mutations and mutations to stop codons. Solid and dashed lines show data from the two replicate fits. For mutations to stop codons, nearly all shift parameters are zero, with a few reaching small non-zero values. For the other mutations, many shift parameters are zero. Of the non-zero parameters, many have small values with only a small fraction reaching large values $>1$. There are more large shift parameters in Delta than BA.2.

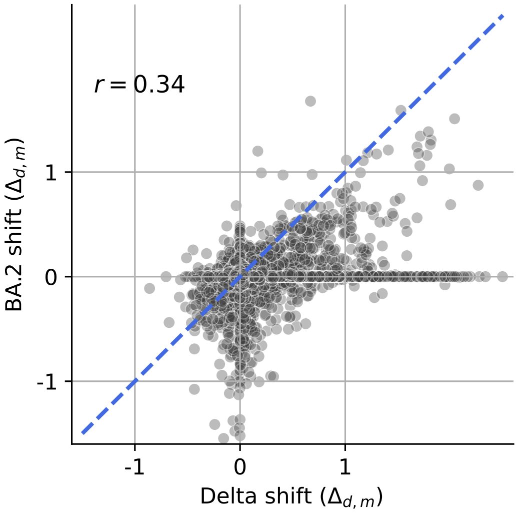


**Fig. S6.** Correlation in the shift parameters inferred for Delta and BA.2, both relative to BA.1. $r$ reports Pearson's correlation coefficient.

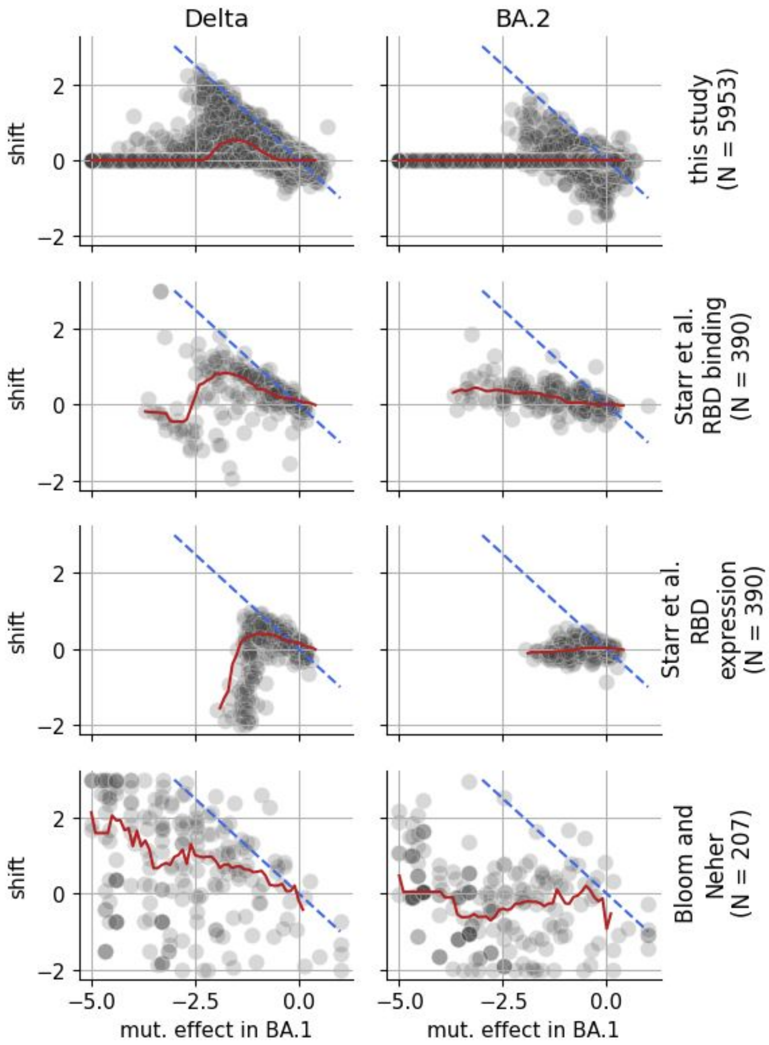


**Fig. S7.** Patterns of mutational effects in BA.1 compared to shifts in Delta or BA.2 in this study and other studies. The top row of plots show data from this study, while the subsequent rows show data from other studies (see row labels on the right). In each row, the left scatter plot shows mutational effects in BA.1 (x-axis) compared to shifts in mutational effects in Delta relative to BA.1 (y-axis), where each dot corresponds to a unique mutation. The right scatter plot shows the same data, but for shifts in mutational effects in BA.2. In each plot, the red line shows the median of all shift parameters within a sliding window of size 1.0 along the x-axis, centered on a given x-value. If a shift occurs along the blue dashed line of $y = -x$, the shift completely neutralizes a mutation's effect. The second and third rows report data from DMS experiments on the RBD measuring the impact of mutations on ACE2 binding, where effects quantify $\log_{10}$ fold change in the $K_d$ of binding, or expression on the surface of yeast, where effects quantify $\log_{10}$ fold change in expression (4, 5). The fourth row reports data from a computational method for estimating mutational effects from millions of naturally occurring sequenced SARS-CoV-2 genomes, with effects independently estimated for each clade (6). For each of these datasets, we computed shifts by subtracting the effect of a given mutation in Delta or BA.2 by its effect in BA.1, only showing data for mutations that overlap between this study and the study in a given row, with row labels indicating the number of mutations being plotted. In this study, shifts in Delta tend to be positive, softening the effects of mutations that are deleterious in BA.1 (top row). As indicated by the red lines, this same qualitative pattern is also visible in the DMS data on RBD-ACE2 binding (second row) and estimates of mutational effects from natural-sequence variation (fourth row), supporting the hypothesis that Delta might be generally more tolerant of mutations than BA.1, though these datasets also contain a subset of mutations with negative shifts, suggesting that this increased mutational tolerance may not be as stark as the shifts from this study suggest. The DMS data on RBD expression (third row) shows a different pattern, where mutations with small deleterious effects in BA.1 tend to be positively shifted in Delta (in agreement with the above trend), though ones with larger deleterious effects in BA.1 tend to have strong negative shifts. This pattern does not support the hypothesis that Delta is more mutationally tolerant than BA.1, at least in terms of mutational effects on RBD expression. In contrast to Delta, the shifts in BA.2 inferred from other datasets do not show strong positive or negative biases as a function of mutational effects in BA.1 on the x-axis, as indicated by the mostly flat red lines at y-values of approximately zero. Note: in the fourth row, data was clipped to the plotted x- and y-ranges to aid with the comparison between studies, though the red lines were not affected by this clipping.

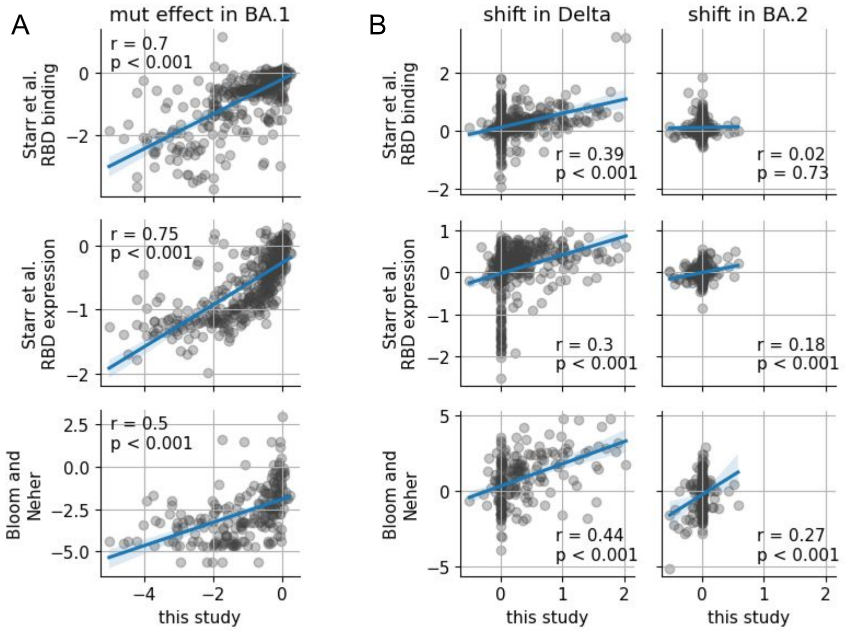


**Fig. S8.** Correlation in mutational effects between this study and other studies. The other studies are the same ones described in Figure S7. **(A)** Each plot compares the mutational effects in BA.1 measured in this study (x-axis) with effects measured from another study (y-axis), where each dot corresponds to a unique mutation present in both studies being compared. Each plot shows an intermediate correlation (r is Pearson's correlation coefficient and p is the corresponding p-value), suggesting that the effects estimated in this study capture biological signal. **(B)** Each plot compares the shifts in mutational effects in either Delta (left column) or BA.2 (right column) measured in this study compared to shifts computed from each dataset in panel A, which we computed here by subtracting the effect of a given mutation in Delta or BA.2 by its effect in BA.1. The shifts in Delta are correlated in all three rows of comparisons, further suggesting that the shifts estimated here capture biological signal, although the the correlations are weak. The shifts in BA.2 show little-to-no correlation with those from the RBD DMSs, perhaps because the largest shifts inferred from our study are outside of the RBD, whereas there is a somewhat higher correlation with shifts estimated from natural-sequence variation of full-length spike, though the overlapping mutations from these datasets still do not include most of the strongly shifted mutations in BA.2 from our study. The reason why these correlations are not higher could be due to noise, as well as differences in selective pressures between datasets (e.g., selection for spike-mediated entry of pseudoviruses into cells is shaped by an overlapping but distinct set of selective pressures compared to selection for ACE2 binding of RBD monomers on the surface of yeast).

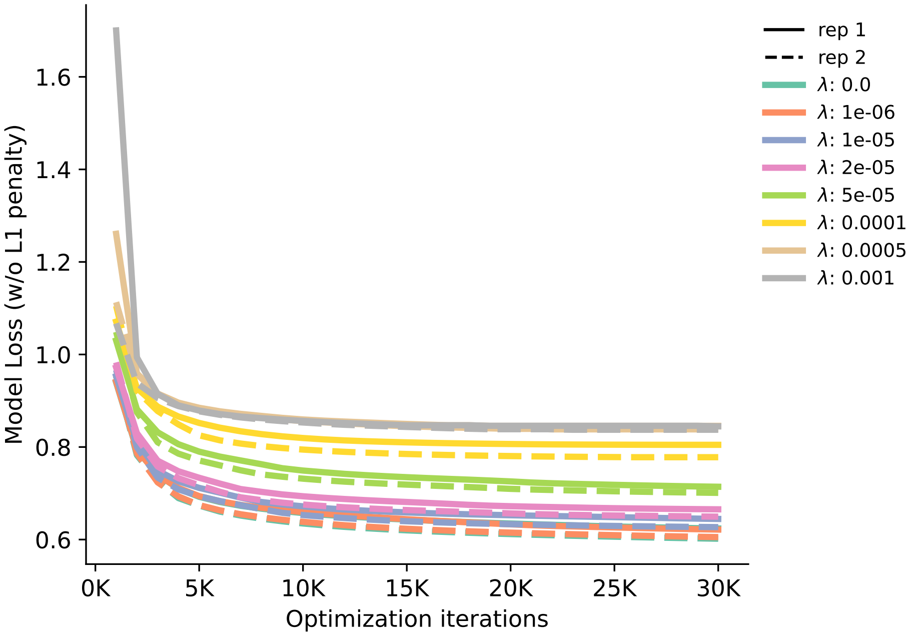


**Fig. S9.** Convergence of model fits at different lasso weights ($\lambda$) and for different biological replicates. We fit all models for 30,000 iterations. The traces show the model loss on the training data (excluding the lasso term) as a function of iteration. The traces approach different asymptotes depending on lasso weight, with replicate fits approaching similar asymptotes.

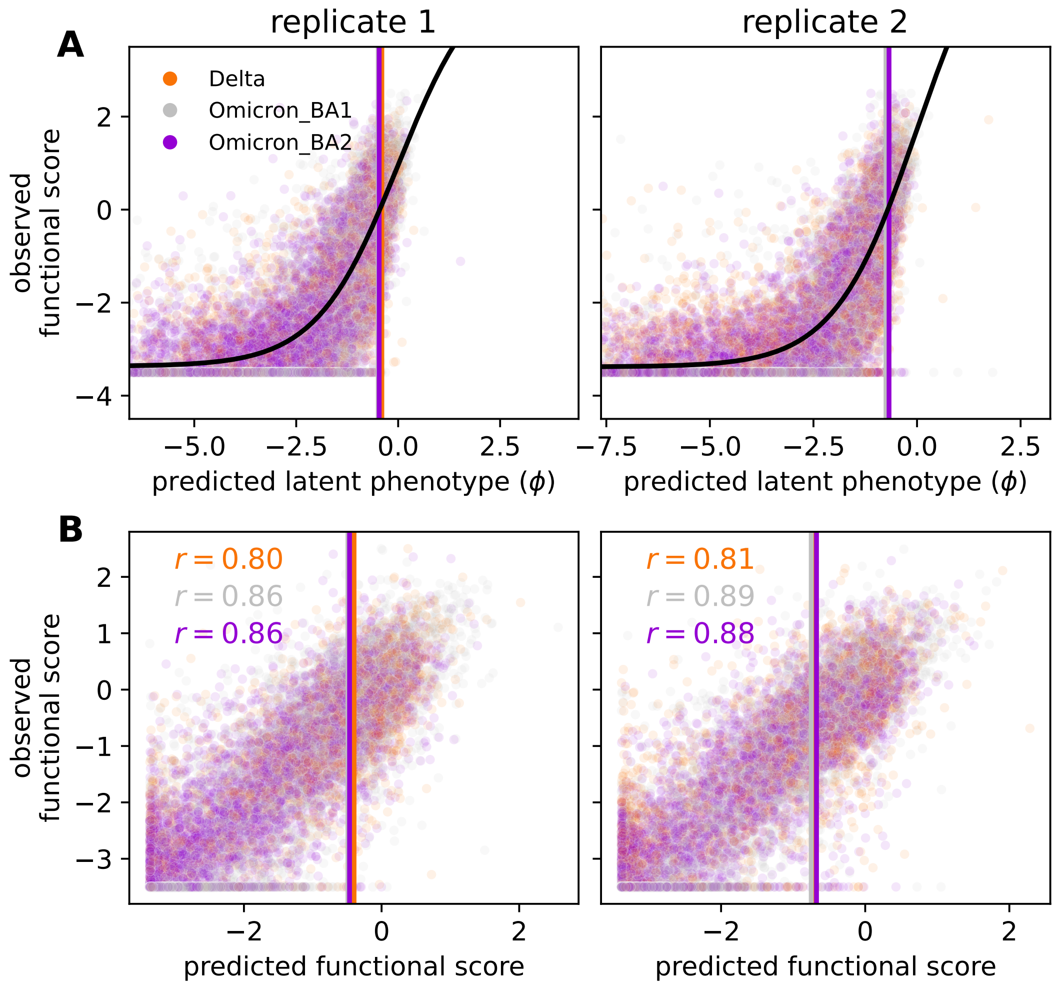


**Fig. S10.** Summary of model fits on training data at a lasso weight of $5 \times 10^{-5}$. The two columns of plots show data for the two replicate fits. $(\mathbf{A})$ In each plot, the black curve shows the inferred global-epistasis function, which is used to map a variant's predicted latent phenotype (x-axis) to a predicted functional score. Each dot corresponds to a single variant from a single experiment, with dots colored by experiment. The y-axis shows the observed functional score of each variant. All dots would be positioned along the sigmoid if predicted and observed functional scores were identical. Instead, the dots track with the sigmoid with some deviations, indicating good fit despite some combination of inaccurate predictions and experimental noise. Vertical lines indicate the inferred latent phenotype of each homolog, all of which cluster together. $(\mathbf{B})$ Correlation between predicted and observed functional scores with dots colored by experiment. Scores were well correlated for each homolog and for each replicate fit, where $r$ reports the corresponding Pearson's correlation coefficient for a given homolog (hue; see legend in panel A).
